# RACER-m Leverages Structural Features for Sparse T Cell Specificity Prediction

**DOI:** 10.1101/2023.08.06.552190

**Authors:** Ailun Wang, Xingcheng Lin, Kevin Ng Chau, José N. Onuchic, Herbert Levine, Jason T. George

**Author notes:** corresponding author: Xingcheng. principal corresponding author August 2023.

## Abstract

Reliable prediction of T cell specificity against antigenic signatures is a formidable task, complicated primarily by the immense diversity of T cell receptor and antigen sequence space and the resulting limited availability of training sets for inferential models. Recent modeling efforts have demonstrated the advantage of incorporating structural information to overcome the need for extensive training sequence data, yet disentangling the heterogeneous TCR-antigen structural interface to accurately predict the MHC-allele-restricted TCR-peptide binding interactions remained challenging. Here, we present RACER-m, a coarse-grained structural template model leveraging key biophysical information from the diversity of publicly available TCR-antigen crystal structures. We find explicit inclusion of structural content substantially reduces the required number of training examples for reliable prediction of TCR-recognition specificity and sensitivity across diverse biological contexts. We demonstrate that our structural model capably identifies biophysically meaningful point-mutants that affect overall binding affinity, distinguishing its ability in predicting TCR specificity of point mutants peptides from alternative sequence-based methods. Collectively, our approach combines biophysical and inferential learning-based methods to predict TCR-peptide binding events using sparse training data. Its application is broadly applicable to studies involving both closely-related and structurally diverse TCR-peptide pairs.

## 1 Introduction

T cell immunity is determined by the interaction of a T cell receptor (TCR) with antigenic peptide (p) presented on the cell surface via major histocompatibility molecules (MHCs) [1]. T cell activation occurs when there is a favorable TCR-pMHC interaction and, for the case of CD8+ effector cells, ultimately results in T cell killing of the pMHC-presenting cell [2]. T cell-mediated antigen recognition confers broad immunity against intracellular pathogens as well as tumor-associated antigenic signatures [3]. Thus, a detailed understanding of the specificity of individual T cells in a repertoire comprised of many (∼ 10^8^) unique T cell clones is required for understanding and accurately predicting many important clinical phenomena, including infection, cancer immunogenicity, and autoimmunity.

Due to the immense combinatorial complexity of antigen (∼ 10^13^) and T cell (∼10^18^) sequence space, initial conceptual process in the field was made by studying simple forms of amino acid interactions, motivated either by protein folding ideas [4, 5] or random energy approaches [6, 7]. Recent advances in high-throughput studies interrogating T cell specificity [8, 9, 10] together with the development of statistical learning approaches have finally enabled data-driven modeling as a tractable approach to this problem. Consequently, a number of approaches have been developed to predict TCR-antigen specificity [11, 12, 13, 14, 15]. A majority of developed approaches input only TCR and pMHC primary sequence information. The persistent challenge with this lies in limited training data given that any reasonable sampling of antigens and T cells, or indeed even an entire human T cell repertoire, represents a very small fraction of sequence space. As a result, many models under-perform on sequences that are moderately dissimilar from their nearest neighbor in the training set, an issue we refer to as *global sparsity*.

While global sparsity complicates inference extension to moderately dissimilar antigens, another distinct challenge exists for reliably predicting the behavior of closely related systems that differ by a single amino acid substitution, which we refer to as *local resolvability*. These ‘point-mutated’ systems require predictive methods capable of quantifying the effects of single amino acid changes on the entire TCR-peptide interaction, a task often limited by lack of sufficient training examples required for reliable estimation of the necessary pairwise residues. Instead, a modeling framework aiming to discern such subtle differences between point-mutants may benefit from learning the general rules of amino acid interactions at the TCR-peptide interface and their varied contributions to binding affinity. Resolving this very particular problem - discerning relevant point-mutations in self-peptide and viral antigens - promises significant therapeutic utility in targeting cancer neoantigens, optimally selecting immune stem cell transplant donors, and predicting the immunological consequences of viral variants. Thus local resolvability represents a distinct learning task wherein detailed reliable pre-dictions need to be made on many small variations around a very specific TCR-pMHC system.

Several structure-based approaches have also been used to better understand TCR-pMHC specificity. Detailed structural models that focus on a comprehensive description of TCR-pMHC interaction, including all-atom simulation and structural relaxation, are computationally limited to describing a few realized systems of interest [16, 17]. Another strategy develops an AlphaFold-based pipeline to generate accurate 3-dimensional structures from primary sequence information to improve the accuracy of TCR-pMHC binding predictions for hundreds of systems [18]. A previous hybrid approach [14] utilized crystal structural data together with known binding sequences to train an optimized binding energy model for describing TCR-pMHC interactions. This approach offered several advantages, including the ability to perform repertoire-level predictions within a reasonable time, along with a reduced demand for extensive training data. However, this model largely focused on a restricted set of peptide or TCR systems using a single MHC-II structural template and did best in explaining mouse I-E^k^-restricted systems. Thus, its ability to make reliable predictions for a structurally diverse collection of TCR and peptide pairs with a conserved human leukocyte antigen (HLA) allele restriction remains unknown.

Here, we leverage all available protein crystal structures of the most common human MHC-I allele variant - HLA-A*02:01 - to develop a combined sequence-structural model of TCR-pMHC specificity that features biophysical information from a diversity of known structural templates. We quantify the structural diversity in available crystal structures of TCR-pMHC complexes[19, 20, 21], and demonstrate that incorporating a small subset of available structural information is sufficient to enable reliable predictions of favorable interactions across a diverse set of TCR-antigen pairs. Our results further suggest that the availability of structural information having close proximity to the true structure of a TCR-pMHC system can ameliorate both global sparsity and local resolvability in discerning the immunogenicity of diverse and point-mutated antigenic variants.

## 2 Results

### Model development and identification of TCR-peptide pairs with structural templates

We build on our previous RACER framework developed primarily on the mouse MHC-II I-E^k^ system [14]. Our new approach, termed RACER multi-template (RACER-m), represents a comprehensive pipeline that leverages published crystal structures of known human TCR-pMHC systems. The training data include every available HLA-A*02:01-restricted system with a published structure [PDB/IEDB] of the TCR-pMHC complex along with their corresponding peptide and TCR variable CDR3*α* and *β* sequences. All associated publications linked to each crystal structure were culled for known strong and weak binding TCR-peptide sequences. Lastly, we included all unique HLA-A*02:01-restricted reads from the ATLAS database [19] comprised of TCR-pMHC systems with reported binding affinity data. In total, 163 unique TCR-peptide pairs and 66 structural templates were identified for training and validation (see Supplementary Data).

We next assessed the structural diversity of training templates by pairwise evaluation of structural similarity using a previously developed method referred to as mutual Q [22, 23]. Mutual Q similarity defines a structural distance metric consisting of a sum of transformed pairwise distances between each residue in two structures normalized within the range of 0 to 1, which was then used to perform hierarchical clustering. We found that the identified structural clusters largely partition TCR-pMHC systems according to immunological function (for example, systems sharing a conserved antigen) with a few exceptions (Fig. 2A). Despite our focus only on a specified HLA-restricted repertoire, the analysis nonetheless revealed significant clustering heterogeneity across all included systems: In some cases (e.g. MART-1, TAX), substantial heterogeneity was observed and associated with significant pairwise dissimilarity of TCR and peptide sequences. This, together with cross-cluster structural diversity, is a consequence of global sparsity given limited observed structures. On the other hand, we also identified structurally homogeneous clusters comprised of TCR-pMHC systems possessing near-identical pairwise sequence similarity (e.g. 1E6), yet these systems have substantial differences in binding affinity, consistent with earlier predictions [6, 7]. This simultaneous manifestation of global sparsity and local resolvability amongst TCR-peptide systems with identical HLA restriction represents a dual challenge for the development of robust predictive models of TCR-peptide specificity.

Given the inter-cluster structural diversity for TCR-pMHC complexes as well as the intra-cluster variability, it is necessary to suitably select a list of structures with sufficient coverage of the identified structural clusters as training data for the model and structural templates for test cases. In particular, we hypothesized that our hybrid structural and sequence-based methodology could benefit from the inclusion of multiple template structures, and the modeling approach presented here was developed with this motivation in mind.

The flow chart in Fig. 1 illustrates the training (top row) and testing (bottom row) algorithm in RACER-m. For training, contact interactions between peptide and TCR were calculated for each of the strong binding systems with available TCR-pMHC crystal structures. Here, contact interactions were defined by a switching function based on the distance between structural residues and a characteristic interaction length (see Methods). For each strong binder, 1000 decoy (weak binder) systems were generated by pairing the original TCR with a randomized version of the peptide. Contact interactions derived from the topology of known TCR-pMHC structures, together with a pairwise 20-by-20 symmetric amino acid energy matrix, determine total binding energy. Each value of the energy matrix corresponds to a particular contribution by an amino acid combination, with negative numbers corresponding to attractive contacts. The training objective aims to select the energy matrix that maximizes separability between the binding energy distributions of strong and weak binders.

**Figure 1:**
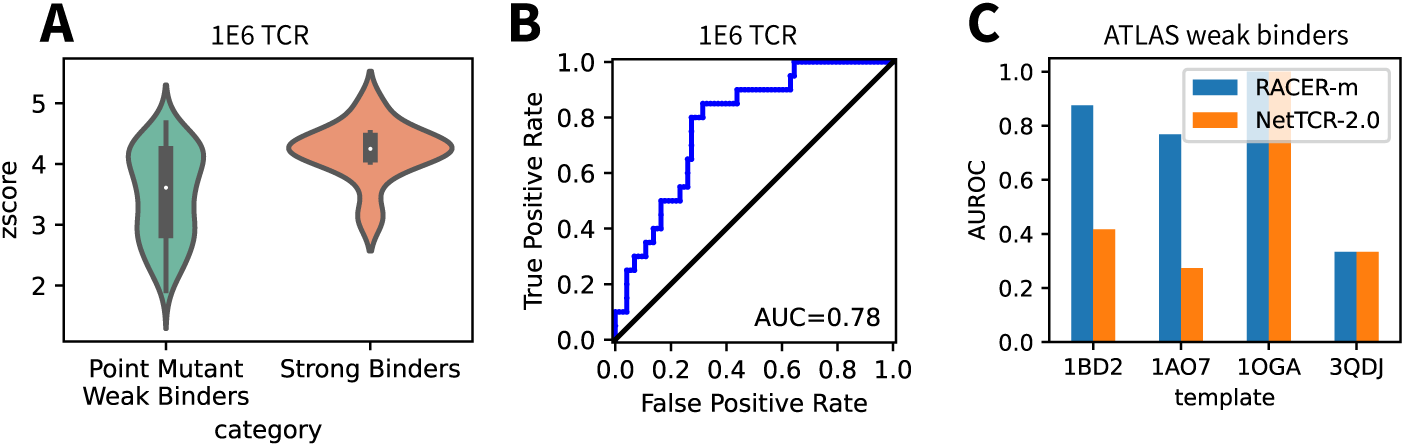
Model architecture of RACER-m. Schematic representation of the training (top row) and testing (bottom row) processes in RACER-m. 66 Crystal structures of known strong binders were used as both training set and template structures for the testing processes, which covers several major clusters of TCR repertoires (MART-1, TAX, 1E6, NLV, FLU) and other clusters with smaller size.

In the testing phase, a sequence threading methods is employed to construct 3D structures for testing cases that lack a solved crystal structure. Here, constructed structures are based on using a chosen known template with shortest (CDR3*α/β* and peptide) sequence distance to the specific testing case. Using the constructed 3D structure, a contact interface can be similarly calculated for each testing case, and 1000 decoy weak binders can be generated by randomizing the peptide sequence. The optimal energy model is then applied to assign energies to the testing system and decoy binders, and the testing system is identified as a strong binder if its predicted binding energy is significantly lower than the decoy energy distributions based on a standardized z score. Here, z score calculation was adopted from the statistical z-test applied to the predicted binding energy of test systems and decoy weak binders, the latter of which were used as a null distribution to compare against a given test binder. The z score of binding energies is defined as *z* = (*E*^̄^_decoy_ − *E*_test_)*/σ*_decoy_, where *E*^̄^_decoy_ is the average predicted binding energy of decoy weak binders, *E*_test_ is the predicted binding energy of the testing system, and *σ*_decoy_ is the standard deviation of the binding energies of decoy weak binders. Testing systems having z scores exceeding 1 are considered strong binding.

### Structural information enhances recognition specificity of pMHC-TCR complexes

RACER-m was developed to explicitly leverage the available structural information obtained from experimentally determined TCR-pMHC complexes for predictions of testing cases. While a prior modeling effort [14] relied on a single structural template for both training and testing and achieved reasonable results given reduced training data, structural differences became prominent as the testing data expanded to include additional TCR and peptide diversity, which resulted in reduced predictive utility. Structural variation has been previously observed and quantified in high molecular detail [24, 25] using docking angles [26] and interface parameters.

For HLA-A*02:01 TCR-pMHC systems, the docking angles^1^ ranged from 29*^◦^* to 73.1*^◦^*, while the incident angle varied from 0.3*^◦^* to 39.5*^◦^* degrees [24, 25, 27]. The observed structural differences among different TCR-pMHC complexes suggest that a single TCR-pMHC complex structure may not accurately represent the contact interfaces of other TCR-pMHC complexes, particularly those with substantially different docking orientations. These distinct docking orientations lead to large variations in the contact interfaces between peptide and CDR3*α*/*β* loops, which can be observed from the diversity in contact maps as shown in Fig. S1. RACER-m overcomes this limitation by the inclusion of 66 TCR-pMHC crystal structures, which are distributed over distinct structural groups, including MART-1, 1E6, TAX, NLV, FLU and serve as both the training dataset and reference template structures for testing cases.

In testing TCR-peptide pairs, all corresponding crystal structures were omitted from predictions. Thus, selecting an appropriate template from available structures became crucial for accurately reconstructing the TCR-pMHC interface and estimating the binding energy. To accomplish this, RACER-m assumed that high sequence similarity corresponds to high similarities in the structure space, which is supported by the correlation between mutual Q score and sequence similarity measured from the 66 solved crystal structures of TCR-pMHC complexes (Fig. S2). This assumption was implemented by calculating sequence similarity scores of the testing peptide and TCR CDR3*α*/*β* sequences with those of all 66 reference templates. In each case, a position-wise uniform hamming distance on amino acid sequences was calculated to quantify the similarity. The sum of CDR3*α* and *β* similarities generated the TCR similarity score, and a composite score was created by taking the product of peptide and TCR scores (see Methods). The template structure having the highest sequence similarity was then selected as the template for threading the sequences of the testing TCR-peptide pair.

To evaluate the extent to which the RACER-m approach can address global sparsity by accurately recapitulating observed specificity in the setting of limited training data, we trained a model using 42.3%^2^ of the total experimentally confirmed strong binders, which sparsely cover all the structural groups involved in the mutual Q analysis shown in Fig. 2A. The remaining 57.7% of TCR-peptide sequences that lack solved structures were utilized as testing cases to validate the sensitivity of the trained energy model. RACER-m effectively recognizes strong binding peptide-TCR pairs and correctly predicts 98.9% of the testing systems using the criteria that z-score is greater than 1. Amongst the 94 testing systems, only one TCR-peptide pair in the TAX structural group was mis-predicted as a weak binders with a binding energy deviating from the average binding energies of decoy weak binders by 0.64*σ*, where *σ* is the standard deviation of the decoy energies. These initial results (Fig. 2) confirm that the model is effectively able to learn the specificity rules from TCR-pMHC systems exhibiting distinct structural representations.

**Figure 2:**
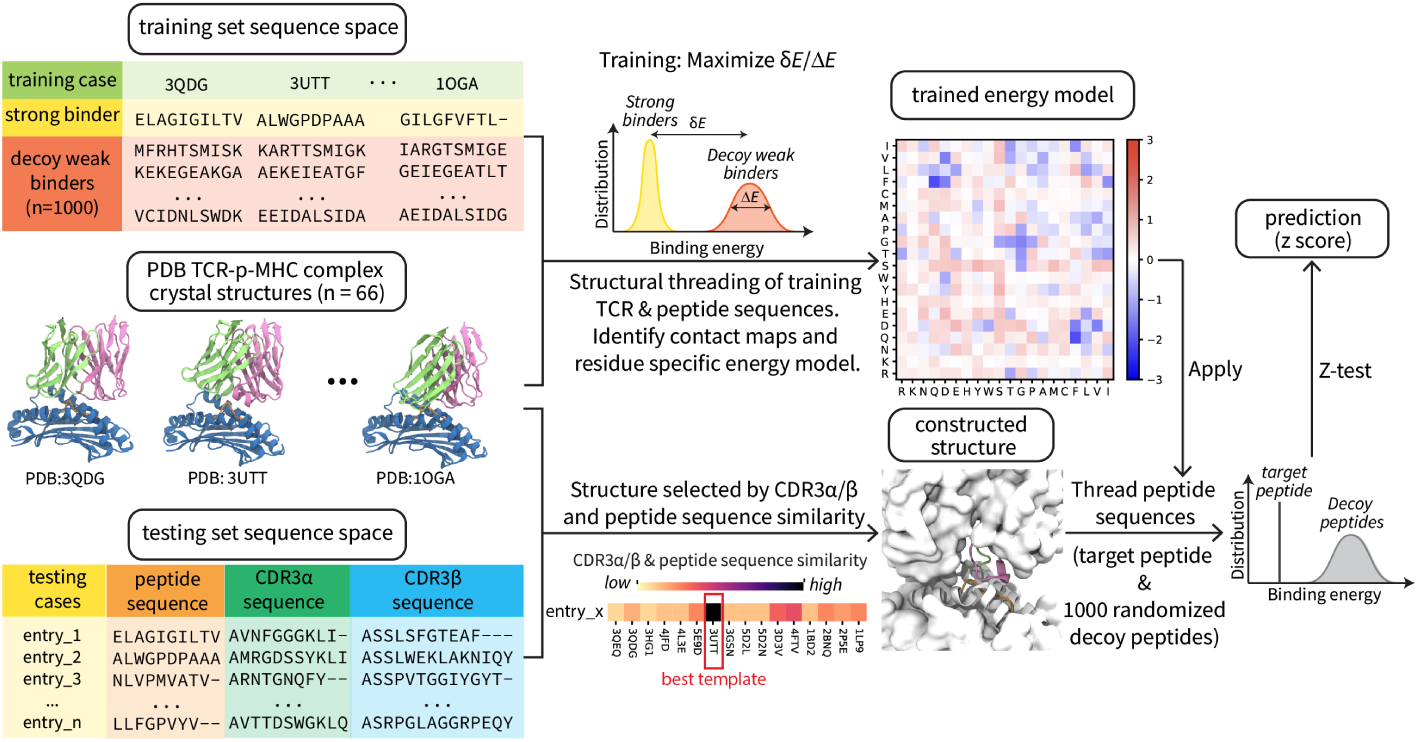
Performance on ATLAS dataset. (A) Mutual Q calculation results between all crystal structures in training set of RACER-m, which measures the structural similarity between every pair of structures from the training set. The linkage map shows the hierarchical clustering result based on the pairwise mutual Q values. Color blocks next to the linkage map indicates the corresponding cluster of the crystal structure in the row. (B) Predicted binding energies for ATLAS dataset (open circles and closed dots) in comparison with the binding energies for corresponding weak binders (box plots). Each open circle represents the predicted binding energy for a structure in the training set, while each closed dot represents the predicted binding energy for a testing case from ATLAS dataset. Each training or testing case is associated with 1000 decoy weak binders generated by randomizing the peptide sequence and pairing with the TCR in the corresponding training/testing structure. Box plots represents the distribution of the predicted energies of the decoy weak bind_1_e_8_rs with the box representing the lower (Q1) to upper (Q3) quartiles and a horizontal line representing the median. The whiskers extended from the box by 1.5IQR, where IQR indicates the interquartile range.

While the reliable identification of strong-binding systems is clinically useful and one important measure of model performance, simultaneous evaluation of model specificity is equally crucial for generating useful predictions on the level of a TCR repertoire. To evaluate the specificity of a global sparsity task, we next tested RACER-m’s ability to discern experimentally confirmed weak-binding systems. We selected peptides or TCRs from the most abundant structural groups (MART-1 and TAX) in the training set to create ‘scrambled’ systems by cross-cluster mismatching of either TCRs or peptides (see Methods for full details). Proceeding in this manner enables a specificity test on biologically realized sequences instead of randomly generated ones. Specifically, every peptide selected from a given structural group (e.g. peptide EAAGIGILTV in the MART-1 group) was mismatched with a list of TCRs specific for peptides belonging to other groups (e.g. TAX, 1E6, FLU, etc.) to form a set of scrambled weak binders.

Following our aforementioned testing protocols, we next calculated z-scores for these mismatched interactions, which were then compared to correctly matched systems with the same peptide sequence (e.g. EAAGIGILTV). We also conducted the complementary test on TCRs using scrambled peptides. The primary advantages of this approach include 1) the ability to match amino acid empirical distributions in binding and non-binding pairs, and 2) utilization of realized TCR sequences for specificity assessment instead of random sequences that possess minimal if any overlap with physiological sequences.

A representative example of these tests utilizing the MART-1 epitope and MART-1-specific TCRs is given in Fig. 3. First, 7 sets of weak binders were constructed by mismatching 36 MART-1-specific TCRs each with 7 non-MART-1 peptides sampled from distinct clusters. We applied RACER-m on each weak binder to predict its binding energy, then compared this value to the distribution of decoy binding energies to obtain a binding z score. z scores of mismatched weak binders, together with those of correctly matched MART-1-TCR strong binders, were used to derive the receiver operating characteristic (ROC) curve (Fig. 3A, Fig. S3). The area under the curve (AUC) was greater than or equal to 0.98 for 5 out of 7 test sets, while the others had AUCs of 0.80 and 0.75, illustrating RACER-m’s ability to successfully distinguish strong binding peptides from mismatched ones in the available MART-1-specific TCR cases.

**Figure 3:**
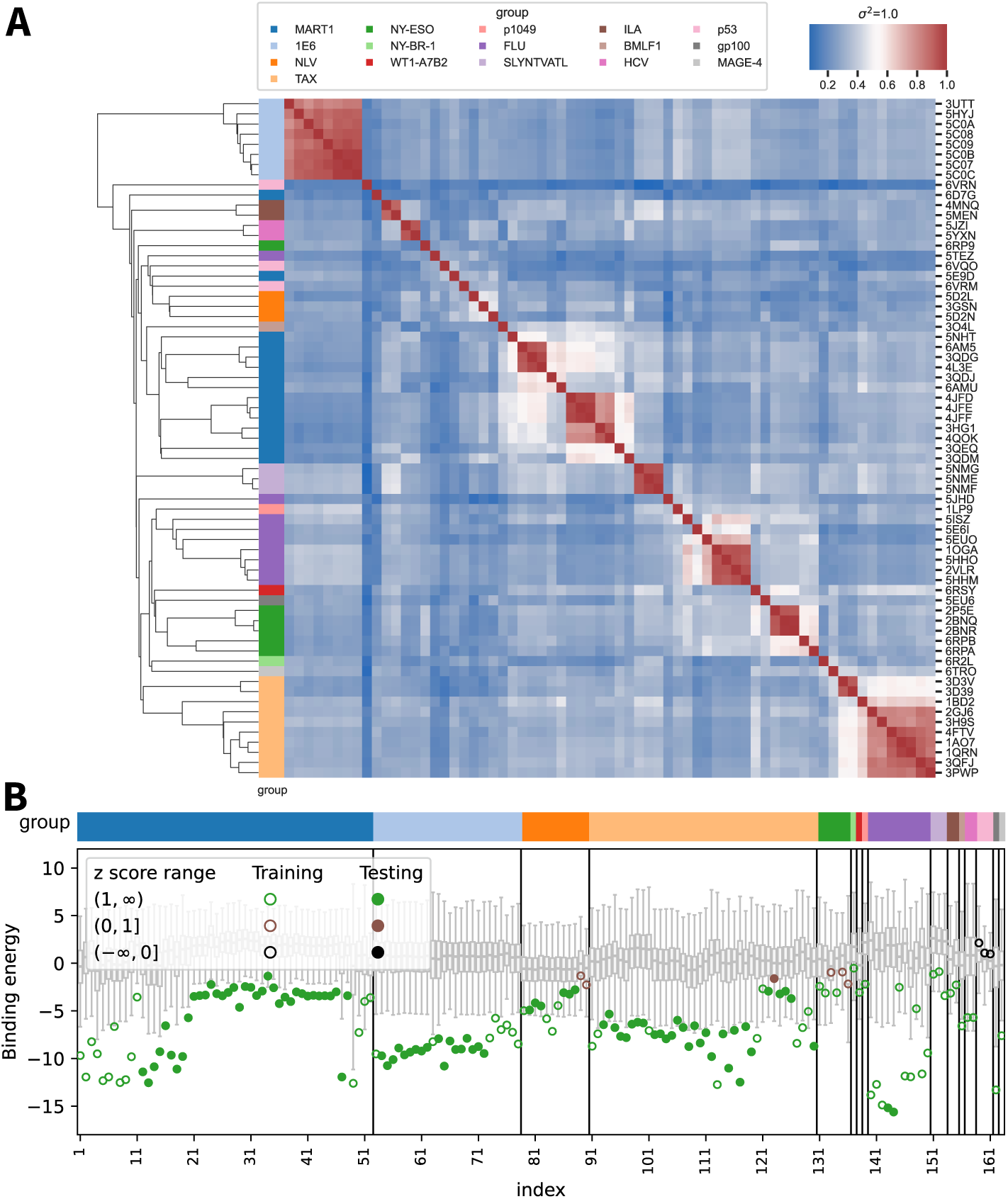
Prediction performance on weak binders generated by mismatching peptides with TCRs. (A) ROC curves for RACER-m classification performance on differentiating weak binders generated by mismatching peptides from NLV, TAX, FLU and 1E6 clusters with MART-1 TCRs from MART-1 strong binders with the same set of TCRs. (B) ROC curves for RACER-m classification performance on distinguishing MART-1 strong binders from mismatched weak binders generated by pairing MART-1 specific peptides with TCRs from NLV, TAX, FLU and 1E6 clusters. (C) When TAX A6 TCR is paired with MART-1 peptide ELAGIGILTV, the Z-score of the mismatched system (triangle) resembles the values from the strong binders (violin shape) formed by the same peptide and TCR RD1-MART1High and its point mutants, which was engineered from A6. In the reverse scenario, TCR RD1-MART1High shows lower Z-score (cross) than TAX strong binders (violin shape) when paired with TAX specific peptide LLFVYPVYV.

An analogous test was performed on the 5 available peptide variants from the MART-1 structural group by mismatching them with 35 TCR sequences contained in the NLV, FLU, 1E6 or TAX clusters. Relative to the binding energies of correctly matched MART-1-specific TCRs, RACER-m performs well in discerning matched vs mismatched TCRs for 4 out of the 5 tested MART-1 peptides (Fig. 3B, Fig. S4), the one initial exception being peptide ELAGIGILTV. Further inspection of the TCRs in this group revealed that the TAX-specific TCR A6 (triangle sign in Fig. 3C) together with several closely associated point mutants had a z score distribution resembling that of the RD1-MART1High TCR and its associated point mutants (Fig. S4E). This could be explained by the fact that the RD1-MART1High TCR was engineered from the A6 TCR to achieve MART-1 specificity [28], wherein A6 was selected because of its similarity with MART-1 specific TCRs in the *V α* region and similar docking mode [28, 29]. However, the engineered (RD1-MART1High) TCR is no longer specific to the TAX peptide (LLFGYPVYV), which is consistent with the z scores predicted from RACER-m. Indeed, when the A6-specific TAX peptide is paired with RD1-MART1High TCR, a relatively lower z score (cross sign in Fig. 3C) is predicted in comparison with the z scores from strong binders (violin shape in Fig. 3C) of the same peptide.

### Evaluation on extended datasets highlights the added value of structural information

Given RACER-m’s performance on the ATLAS data, we then applied the model to additional datasets to further validate its ability in the setting of global sparsity. The 10x genomics [30] dataset details many TCR-peptide binders collected from five healthy donors. HLA-A*02:01-restricted samples in this dataset include 23 unique peptides, and the number of TCRs specific for each peptide varied from 8365 (e.g. GILGFVFTL) to 1 (e.g. ILKEPVHGV). We remark that the diversity of HLA-A*02:01 samples was significantly reduced to 1741 systems having unique CDR3*α*/*β* and peptide sequences after removing redundancies. Importantly, we selected this large dataset as a reasonable test since 89.26% of the 1741 testing systems did not share either the same CDR3*α* or CDR3*β* sequence in common with the list of available systems used in the training set, and 99.89% of the testing systems did not have the same CDR3*α*-CDR3*β* combination with the training set, although 7 out of the 23 peptides were shared with the training set.

Given this relative lack of overlap with our training data, we applied RACER-m to all unique HLA-A*02:01 pairs. In a majority (88.9%) of these cases across a large immunological diversity of peptides, RACER-m successfully identifies enriched z scores in the distribution of binding TCRs (Fig. 4A). The distinction of TCRs belonging to testing vs. training sets, together with the striking difference in the size of training and testing systems, suggest that shared structural features were able to augment RACER-m’s predictive power on distinct tests. Thus, the inclusion of structural information in model training enhances RACER-m’s predictive ability across distinct TCR-pMHC tests. There were several cases where RACER-m’s predicted distributions overlapped significantly with low z scores, indicating a failed prediction; in these cases we investigated whether this could be explained by the lack of an appropriate structural template. A significant positive correlation was observed between a testing case’s optimal structural template similarity and the RACER-m-predicted z scores, consistent with a decline in model applicability whenever the closest available template is inadequate for representing the system in question (Fig. S5). Despite this, the RACER-m approach, trained on 69 cases, was able to predict roughly 90% of strong binders contained in over 1700 distinct testing cases in the 10x genomics dataset.

**Figure 4:**
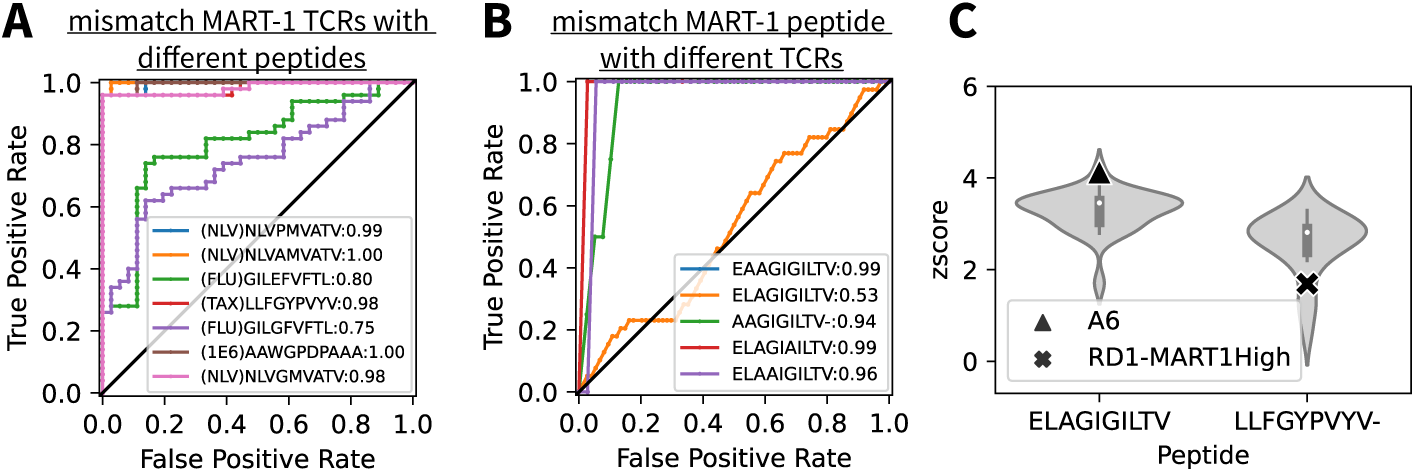
Validate the predictive power of RACER-m with external datasets. (A) Prediction results of RACER-m on the HLA-A*02:01 restricted systems from 10x Genomics dataset collected from 5 healthy donors. 1741 unique pairs of TCR-peptide sequences were tested and the prediction results of z score were grouped by the immunological profile of the test systems and depicted as box plots. (B) Comparison of classification performance between RACER-m and NetTCR-2.0 on a curated list of public TCR-pMHC repertoires [12] comprised by both strong binders and mismatched weak binder. Due to the restriction of NetTCR-2.0 on the peptide length (9-mer), there is no data from NetTCR-2.0 for the two 10-mer peptides (KLVALGINAV and ELAGIGILTV), (C) The classification performance of RACER-m on another set of TCR-pMHC test systems [31].

We then compared RACER-m’s performance to NetTCR-2.0 [11], a well-established convolutional neural network model for predictions of TCR-peptide binding that is trained on over 16000 combinations of peptide/CDR3*α*/*β* sequences. This comparison was performed on a publicly available list of TCR-pMHC repertoires curated by Zhang *et al.* [12] which were mutually independent of RACER-m or NetTCR-2.0 training data, wherein we included known strong binders and mismatched weak binders for 8 unique peptides of HLA-A*02:01. Since NetTCR-2.0 has a restricted length for antigen peptide (no longer than 9-mer), it cannot be applied on testing systems with 10-mer peptides such as KLVALGINAV and ELAGIGILTV, which are absent from the NetTCR-2.0 evaluation in Fig. 4B. The area under the ROC curve (AUROC) was used as a standard measure of classification success. In the majority of cases, RACERm outperformed NetTCR-2.0 in diagnostic accuracy with higher ROC values (Fig. 4B). Lastly, RACER-m was further evaluated using an unrelated set of TCR-pMHC data comprised of 400 samples made up of the strong binders and mismatched weak binders with 4 peptides and 100 TCRs [31], which also gives us good distributional performance (Fig. 4C). In one of the 4 peptides included in this dataset, RACER-m seems to have difficulty providing correct classification about strong and weak binders for peptide CVNGSCFTV, which could again be explained by the lack of appropriate structure templates for this pMHC and related strong binding TCRs (Fig. S6).

### RACER-m specificity of point-mutated variants and preservation of local resolvability

Encouraged by model handling of global sparsity in tests of disparate binding systems having high sequence diversity, we next evaluated RACER-m’s ability in maintaining local resolvability of point-mutated peptides with near-identical sequence similarity to a known strong binder, which represents a distinct and usually more difficult computational problem. Understanding in detail which available point mutants enhance or break immunogenicity is directly relevant for assessing the efficacy of tumor neoantigens and T cell responses to viral evolution. Additionally, the performance of structural models in accomplishing this task are a direct readout on their utility over sequencebased methods, since the latter case will struggle to accurately cluster, and therefore resolve, systems having single amino acid differences. To evaluate RACER-m’s ability to recognize point mutants, we performed an additional test on an independent comprehensive dataset of TCR 1E6 containing a point mutagenic screening of the peptide displayed on MHC. This testing set includes 20 strong binders and 73 weak binders [21], wherein strong binding to the 1E6 TCR was confirmed by TNF*α* activity. RACER-m demonstrates enrichment of the distribution of binding energies for strong binders vs. confirmed weak cases (Fig. 5A). ROC analysis of the RACER-m’s ability to resolve these groups gives an AUC of 0.78. It is important to note that only 2 strong binders of this system were included in the training of RACER-m’s energy model.

**Figure 5:**
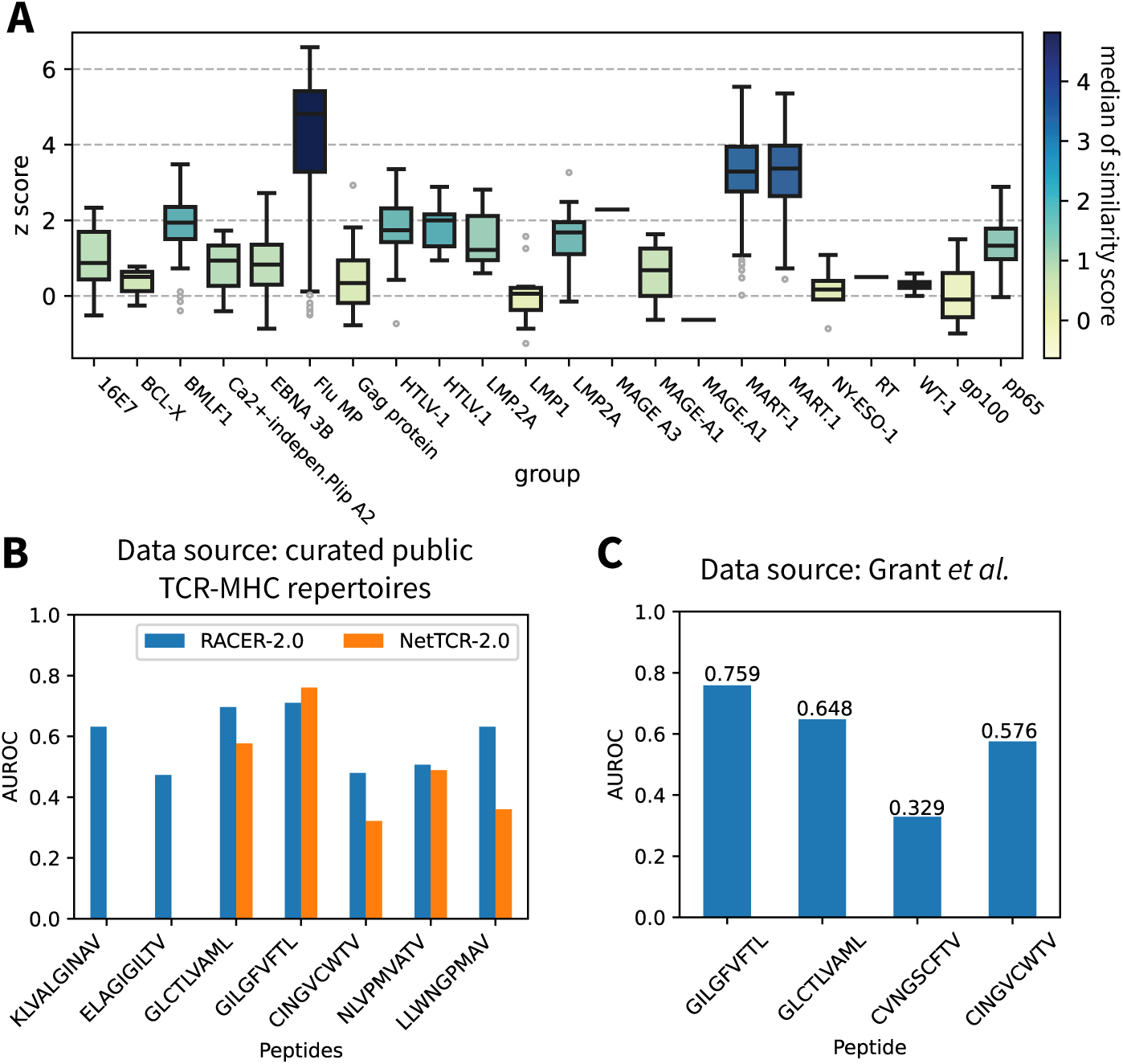
RACER-m’s performance on differentiating strong binders from pointmutant weak binders. (A) Distribution of z scores from strong binders of 1E6 TCR and weak binders from point mutagenic screen. (B) ROC curve for RACER-m classification performance using the strong and point-mutant weak binders for 1E6 TCR. (C) Comparison of RACER-m and NetTCR-2.0 in classification of strong and point-mutant weak binders from ATLAS dataset.

Inspired by these initial results on the 1E6 mutagenic screen, we extended this analysis to all point-mutated weak binding systems in the ATLAS dataset, specifically those with *K_D_* values greater than 200 *µ*M. Our results, presented template-wise for each structure in the point-mutant data, demonstrate that RACER-m improves in this recognition task when compared to NetTCR-2.0 (Fig. 5C). Lastly, to explicitly explore the strength of structural modeling in predicting the impact of small but immunologically significant single amino acid differences, we quantified the predicted z scores for both strong and weak binders as a function of sequence similarity (Fig. S7). The results demonstrate that the inclusion of information from correctly identified structural templates enhances RACER-m’s predictive power. Collectively, our results suggest that RACER-m offers a unique computational advantage over traditional, sequenceonly methods of prediction by leveraging significantly fewer training sequences with key structural information to efficiently identify the contribution of each amino acid change.

## 3 Discussion

Reliable and efficient estimation of TCR-pMHC interactions is of central importance in understanding, and thus optimizing, the adaptive immune response. Decoding the predictive rules of TCR-pMHC specificity is a formidable challenge, largely owing to the extreme sparsity of available training data relative to the diversity of sequences that need to be interrogated in meaningful investigation. We developed RACER-m to augment the predictive power of a relatively small number of TCR and epitope sequences by leveraging the structural information contained in solved TCR-pMHC crystal structures. Our analysis focused on the most common human MHC allele variant, due to the abundance of sequence and structural data. Despite this restriction, we observed structural heterogeneity underpinning the specificity of various TCR-pMHC systems in distinct immunological contexts. Enhancement in predictive accuracy was largely driven by the availability of a small list of structural templates, which included 66 crystal structures of TCR-pMHC complexes from the Protein Data Bank.

Using our minimal list, together with mutually independent testing systems for RACER-m and NetTCR-2.0, we find that our model is able to outperform on both detection of strong binders as well as avoidance of weak binders - both representing distinct but equally important tasks. We advocate for the inclusion of such mixed performative tests for rigorous validation as a necessary and standardized component in model evaluation, in addition to model comparisons using testing data that is equally dissimilar from the training data included in competing models.

Intriguingly, incorporation of structural information into the training approach enables the development of a model that maintains predictive accuracy while dealing with both global sparsity and local resolvability, all while requiring substantially reduced training sequence data. Our results suggest that a wealth of information is contained in the structural templates pertaining to key contributors of a favorable TCR-peptide interaction, wherein conserved features across distinct systems can be learned to mitigate global sparsity. Conversely, structural encoding of information pertinent to residues whose amino acid substitutions either preserve or break immunogenicity also assists RACER-m trained on only a small subset of all possible point-mutagens by identifying key contributing positions and residues, thereby preserving local resolvability.

Moreover, model accuracy correlated directly with the availability of a template having sufficient proximity to the sequences of testing systems. As a result, we anticipate that RACER-m will improve as more structures become readily available for inclusion. Existing computational methods for identifying structural models from primary sequence data [18] may provide an efficient method of adding highly informative structures into the candidate pool for testing. This, together with identifying the minimal sufficient number of distinct structural classes within a given MHC allele restriction remain tasks for subsequent investigation. Our current results suggest this is doable given the small number of structures available for explaining the diverse systems studied herein. Significantly, the inclusion of only 66 template structure augmented RACER-m’s ability to accurately differentiate strong and weak binders when evaluated with hundreds and even thousands of testing systems. This structural advantage was enhanced both by the approach of hybridizing sequence and structural information into the training and testing protocols and the availability of templates that shared sufficient sequence-based similarity to testing cases so that an adequate threading template was available.

## 4 Methods

### RACER-m Model

To predict the binding affinity between a given TCR-peptide pair, we employed a pairwise energy model to assess the TCR-peptide binding energy [14]. The CDR3*α* & CDR3*β* regions were used to differentiate between different TCRs because CDR3 loops primarily interact with the antigen peptides while CDR1 and CDR2 interact with MHC [32]. However, the binding energy was evaluated based on the entire binding interface between TCR and peptide. As illustrated in Fig. 1, we included 66 experimentally determined TCR-p-MHC complex structures and 3 additional TCR-p-MHC complex structures composed of experimentally determined p-MHC complexes with corresponding TCR structures as strong binders for training an energy model (details in Supporting Methods), which was subsequently used to evaluate binding energies of other TCR-peptide pairs based on their CDR3 and peptide sequences. Additionally, for each strong binder, we generated 1000 decoy binders by randomizing the peptide sequence. These 69,000 decoys constitute an ensemble of weak binders within our training set.

To parameterize this energy model, we optimized the parameters by maximizing the gap of binding energies between the strong and weak TCR-peptide binders, represented by *δE* in Fig. 1. The resulting optimized energy model will be used for predicting the binding specificity of a peptide towards a given TCR based on their sequences. Further details regarding the calculation of binding energy are provided below.

### Detailed calculation of TCR-peptide binding energies

To evaluate the binding affinity between a TCR and a peptide, RACER-m utilized the framework of the AWSEM force field [33], which is a residue-resolution protein force field widely used for studying protein folding and binding [33, 34]. To adapt the AWSEM force field for predicting TCR-peptide binding energy, we utilized its direct protein-protein interaction component to calculate the inter-residue contacting interactions at the TCR-peptide interface. Specifically, we utilized the C*β* atoms (except for glycine, where C*α* atom was used instead) of each residue to calculate the contacting energy using the following expression:

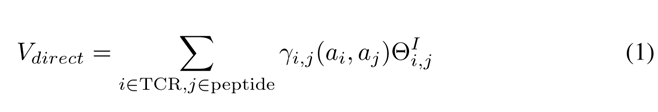

In Eq. 1, Θ*_i,j_* represents a switching function that defines the effective range of interactions between each amino acid from the peptide and the TCR:

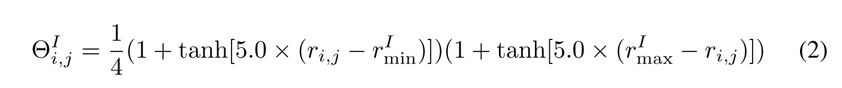

where *r^I^* = 6.5Å and *r^I^* = 8.5Å. The coefficients *γ_i,j_*(*a_i_, a_j_*) define the strength of interactions based on the types of amino acids (*a_i_, a_j_*). The *γ_i,j_*(*a_i_, a_j_*) coefficients are also the parameters that are trained in the optimization protocols described as follows.

### Optimization of energy model for predicting the TCR-peptide binding specificity

To predict the binding specificity between a given TCR and peptide, the energy model is trained using interactions gathered from the known strong binders and their corresponding randomly generated decoy binders. Following the protocol specified in our previous paper [14], the energy model of RACER-m was trained to maximize the gap between the binding energies of strong and weak binders. In addition, a larger training set was used to achieve a more comprehensive coverage of the structural and sequence space. Specifically, the binding energies were calculated from individual strong binders (*E*_strong_) and their corresponding decoy weak binders (*E*_decoy_) as described in Eq. 1. We then calculated the average binding energy of the strong (⟨*E*_strong_⟩), the average binding energy of the decoy weak binders (⟨*E*_decoy_⟩), and the standard deviation of the energies of the decoy weak binders (Δ*E*).

To train the model, the parameters *γ_i,j_*(*a_i_, a_i_*) were optimized to maximize *δE/*Δ*E*, where *δE* = ⟨*E*_decoy_⟩ − ⟨*E*_strong_⟩, resulting in the maximal separation between strong and weak binders. Mathematically, *δE* can be represented as **A**^⊺^*γ*, where

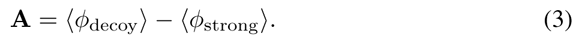

Furthermore, the standard deviation of the decoy binding energies Δ*E* can be calculated as Δ*E*^2^ = *γ*^⊺^*Bγ*, where

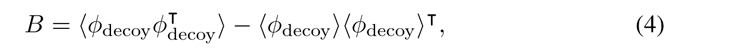

here, *ϕ* takes the functional form of *V_direct_* and summarizes interactions between different types of amino acids. Therefore, the vector **A** specifies the difference in interaction strengths for each pair of amino acid types between the strong and decoy binders, with a dimension of (1,210), while the matrix *B* is a covariance matrix with a dimension of (210, 210).

With the definition above, maximizing the objective function of *δE/*Δ*E* can be reformulated as maximization of **A**^⊺^*γ/*^√^*γ*^⊺^*Bγ*. This maximization can be effectively achieved through maximizing the functional objective *R*(*γ*) = A^⊺^*γ* − *λ*_1_^√^*γ*^⊺^*Bγ*. By setting *∂R*(*γ*)*/∂γ*^⊺^ to 0, the optimization process leads to *γ B^−^*^1^**A**, where *γ* is a (210, 1) vector encoding the trained strength of each type of amino acid-amino acid interactions. For visualization purposes, the vector *γ* is reshaped into a symmetric 20-by-20 matrix, as shown in Fig.1. Additionally, a filter is applied to reduce the noise caused by the finite sampling of decoy binders. In this filter, the first 50 eigenvalues of the *B* matrix are retained, and the remaining eigenvalues are replaced with the 50^th^ eigenvalue.

### Construction of target TCR-p-MHC complex structures from sequences

Since RACER-m calculates the binding energy based on the interaction contacts between a given peptide and a TCR, it relies on the 3D structure of the TCR-p-MHC complex for contact calculation. Although the training data include a 3D structure for each of the TCR-peptide strong binders, we usually lack 3D structures for most of the testing cases. To address this limitation, we used the software Modeller [35] to construct a structure based on the target peptide/CDR3 sequences in the test system and a template crystal structure selected from the training set.

Specifically, for each testing system, a position-wise uniform Hamming distance was computed between the target sequence and each of the sequences from the 66 training strong binders with complete TCR-p-MHC complex structures, separately for peptide, CDR3*α*, and CDR3*β* regions. Then, sequence similarity scores were assigned to peptide, CDR3*α*, and CDR3*β*, respectively with the number of amino acids that remain the same between target and template sequences. To calculate a composite similarity score for the target TCR-peptide complex, we summed the similarity scores of the CDR3*α* and *β* regions and multiplied this sum by the peptide similarity score. The template structure with the highest similarity score was selected as the template for the subsequent sequence replacement using Modeller (Fig. 1 bottom).

To perform the sequence replacement, the peptide, CDR3*α*, and CDR3*β* sequences in the template structure were replaced with the corresponding target sequences in the testing TCR-peptide system. The location of the target sequence in the template structure was determined by aligning the first amino acid of the target sequence with the original template sequence. If the two sequences had different lengths, the remaining locations were patched with gaps. This sequence alignment and the selected template structure were then used as input for Modeller to generate a new structure. The constructed structure was then used for the estimation of the binding energy of the testing system.

### Generation of weak binders by mismatching sequences of known TCR-peptide pairs

To test the performance of RACER-m in distinguishing strongly bound TCR-peptide pairs from weak binders, we generated a set of weak binders by introducing sequence mismatches between the peptides and TCRs from the known strongly bound TCR-peptide pairs. As shown in Fig. 2, the strong binders were grouped based on their immunological systems, such as MART-1 and TAX. It is important to note that pairs within the same group also share similar TCR-peptide structural interfaces.

To generate the weak binders, we mismatched the sequences of peptides and the CDR3*α*/*β* pairs from different groups. For example, 36 pairs of MART-1 specific CDR3*α*/*β* sequences were mismatched with 7 non-MART-1 peptides to form weak binders for Fig. 3A, while 5 MART-1 specific peptides were mismatched with 35 pairs of non-MART-1 CDR3*α*/*β* sequences to form weak binders in Fig. 3B. The newly generated combinations of sequences were then used to create 3D structures of the TCR-p-MHC complexes, following the protocol specified in Section *Constructing TCR-p-MHC complex structure from sequence*.

### Mutual Q calculation

To quantify the structural distances between the 66 crystal structures of TCR-p-MHC complexes, a pairwise mutual Q score was used to calculate the structural similarity between every pair of the 66 structures. Since our focus is on the contact interface between the peptide and the CDR3*α*/CDR3*β* loops of the TCR, the mutual Q score was computed between these regions. We adopted a similar protocol used in [22] and calculated the mutual Q score between structures A and B with the following expression:

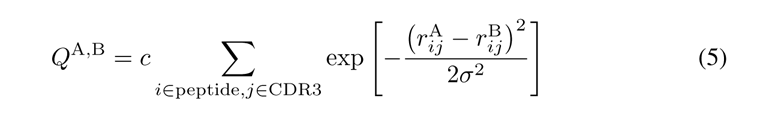

where *i* and *j* are indices of atoms from the peptide and CDR3 loops, respectively. *r*^A^_*ij*_ and *r*^B^_*ij*_ denote the contact distances between atom *i* and *j* in structure A and B respectively. For simplicity, *σ* was set as 1 Å instead of using the sequence distance between *i* and *j* as done in [22]. The coefficient *c* normalizes the value of *Q* to fall within the range of 0 and 1. This definition ensures that a larger value of *Q* indicates a greater structural similarity between the two systems.

### Prediction protocols with NetTCR-2.0

To test the predictive performance of RACER-m, we compared the prediction accuracy of RACER-m with NetTCR-2.0, another widely used computational tool trained with a convolutional neural network model, as described by Montemurro *et al.* [11]. To ensure a fair comparison, we retrained the NetTCR-2.0 model with the paired alpha beta dataset with a 95% partitioning threshold (file train ab 95 alphabeta.csv, provided in https://github.com/mnielLab/NetTCR-2.0). The trained model was then used to classify the strong and weak binders, as shown in Fig. 5C. Due to the peptide length restriction in the application of NetTCR-2.0, we excluded peptides longer than 9 residues from our testing prediction.

## Acknowledgments

Work by the Center for Theoretical Biological Physics was supported by the NSF (Grant PHY-2019745). JTG was supported by CPRIT grant RR210080. JNO was also supported by the NSF grant PHY-2210291 and by the Welch Foundation (Grant C-1792). JNO and JTG are CPRIT Scholars in Cancer Research.

## Footnotes

1 The docking angle is the angle between the peptide binding groove on the MHC and the vector between the TCR domains, the latter is calculated using th centroids of the conserved disulfide bonds in each domain. This angle corresponds to the twist of the TCR over the p-MHC.

2 In addition to the 66 crystal structures of HLA-A*02:01 TCR-pMHC systems, 3 strong binders (PDB: 3GSR, 3GSU, and 3GSV) of NLV peptide with solved pMHC structures were also included in the training set. See Supporting Methods for details.

## Notes

### Competing Interest Statement

The authors have declared no competing interest.

