## Supplementary Information for "RACER-m Leverages Structural Features for Sparse T Cell Specificity Prediction"

August 2023

### Contents

**1 Supporting Methods** **3**

1.1 Training data selection for RACER-m. . . . . . . . . . . . . . . . . . . . . . . . . . 3

1.2 Collection of point-mutant weak binders for 1E6. . . . . . . . . . . . . . . . . . . . . 3

**2 Supporting Figures** **3**

### 1 Supporting Methods

#### 1.1 Training data selection for RACER-m.

The RACER-m training set consists of TCR-p-MHC complex structures restricted to the HLA-A*02:01 allele, collected from the Protein Data Bank, which initially comprises 66 complex structures. However, it was observed that when trained on these 66 structures, RACER-m systematically underestimated the binding affinities of strong binders specific to NLV peptides and their variants. To address this issue, we incorporated three additional structures of strong binders from [1]in which 6 strong binders were reported when combining NLV variants with TCR RA14 and 3 of them were provided with p-MHC structures. By combining these p-MHC structures with TCR RA14 to form the TCR-p-MHC complex structure and adding them as supplementary training cases, we expanded the training set to a size of 69. The inclusion of these three NLV strong binders effectively resolved the systematic underestimation problem concerning the predictions of NLV-specific strong binders, while preserving the excellent predictive power for other strong binders in the ATLAS dataset.

#### 1.2 Collection of point-mutant weak binders for 1E6.

To test the performance of RACER-m in terms of discerning strong binders from point-mutant weak binders, we collected point-mutant weak binders from a comprehensive peptide-mutagenesis study by Bulek *et al.* [2]. Through the mutational scan, Bulek *et al.* assessed the impact of point-mutations on the binding of peptide ALWGPDPAAA to the 1E6 TCR with the tumor necrosis factor (TNF). Since it was pointed out that the 1E6 TCR was tolerant to changes in peptide residues Ala1, Leu2, Ala8, Ala9 and Ala10 [2], we collected point-mutations at positions 3 to 7 with TNF equal or smaller than 25, and considered them as point-mutant weak binders for 1E6.

### 2 Supporting Figures

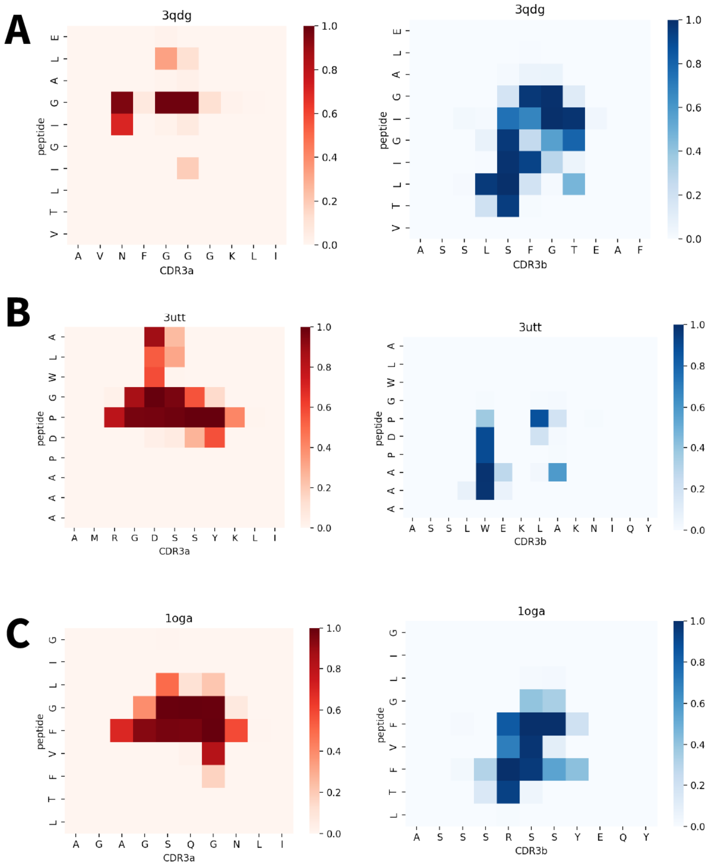


Figure S1: **Contact maps of crystal structures 3QDG, 3UTT and 1OGA.** Each contact map was calculated by measuring the proximity $W_{i,j}$ between each residues of peptide (residue $i$) and CDR loops (residue $j$) based on their mutual distance ($d$) using a smoothed step function: $W_{i,j} = (1 - \tanh(d - d_{max}))/2$, where $d_{max} = 8.5$Å. Only C$\beta$ atoms were used for the mutual distance calculation (except for glycine, where the C$\alpha$ atom was used).

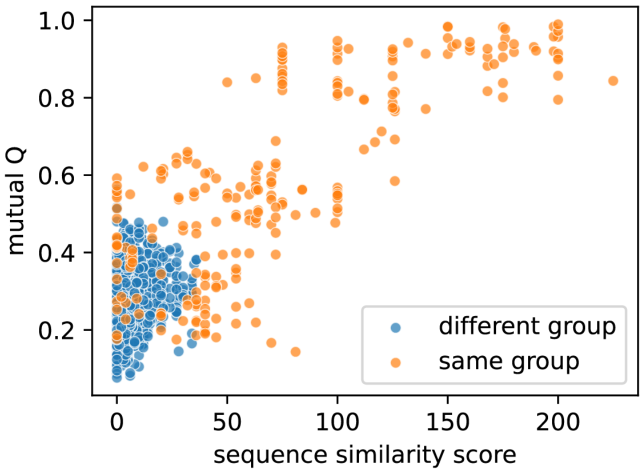


Figure S2: **Relationship between structure and sequence similarities of TCR-pMHC complexes.**

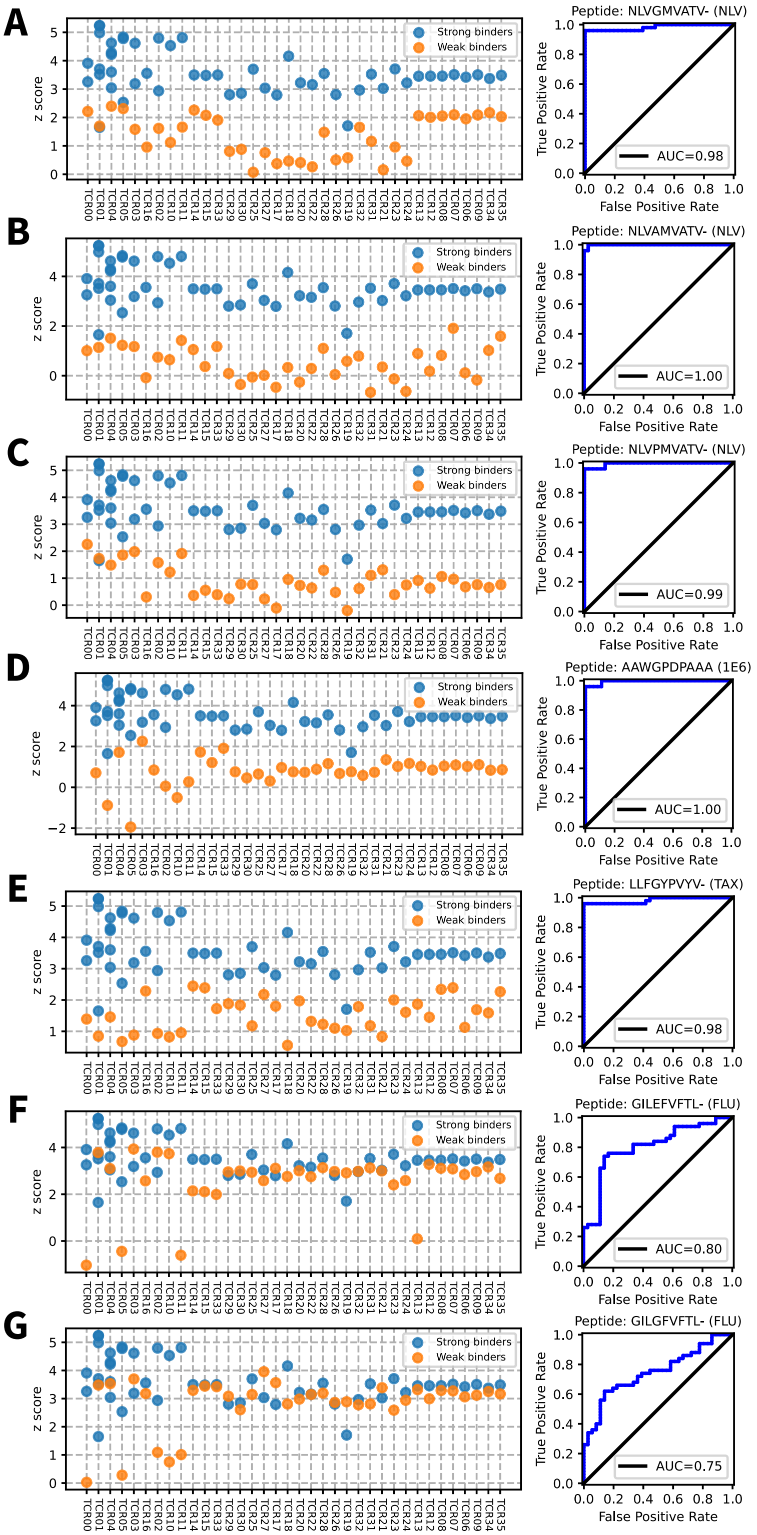




Figure S3: **Comparison of predicted z-scores between strong binders of MART-1 (blue) and weak binders (orange) generated by mismatching MART-1 TCRs with peptides from 1E6, TAX, NLV, and FLU.**

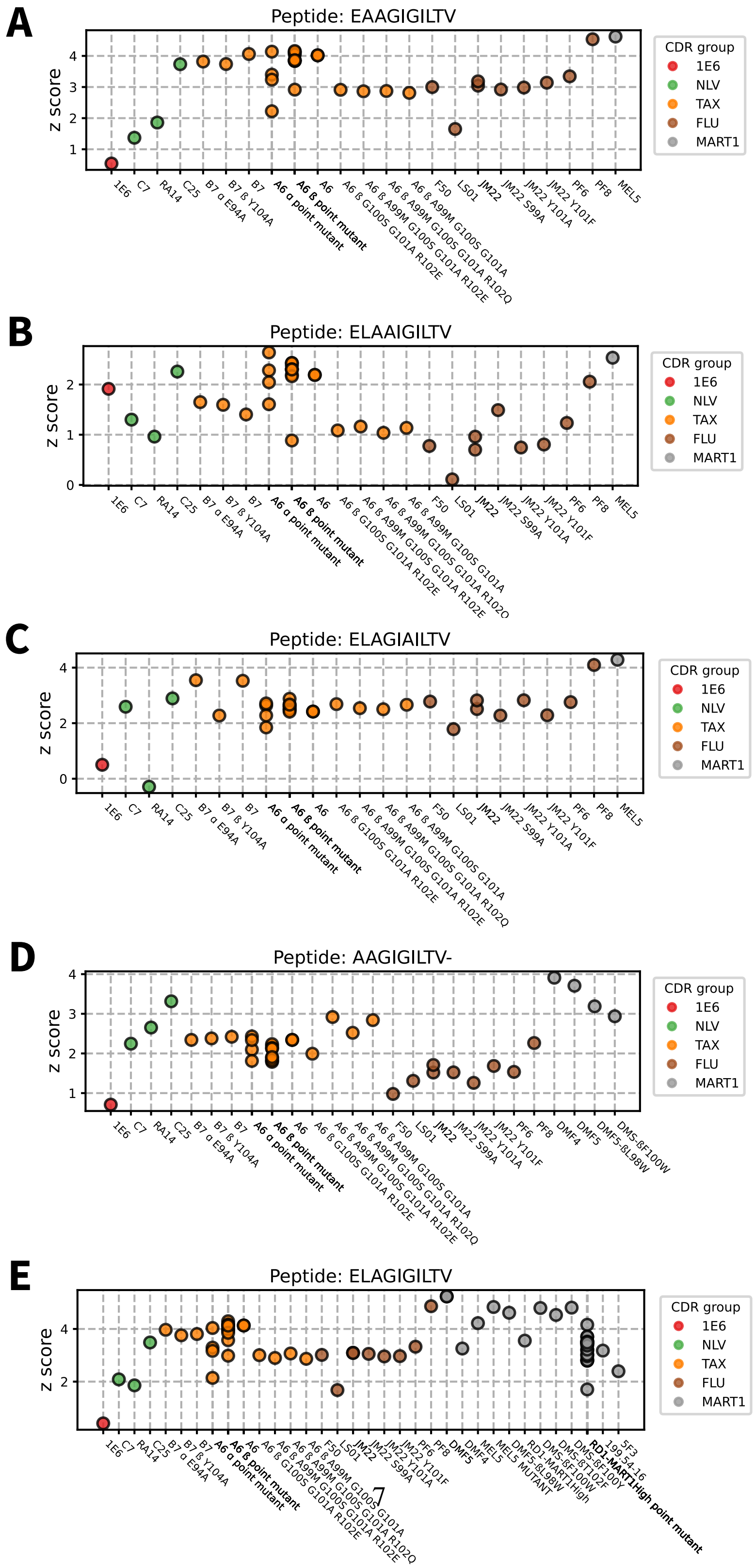


Figure S4: **Comparison of predicted z-scores between strong binders of MART-1 (grey) and weak binders (red, green, orange, and brown) generated by mismatching MART-1 peptides with TCRs specific to 1E6, TAX, NLV, and FLU.**

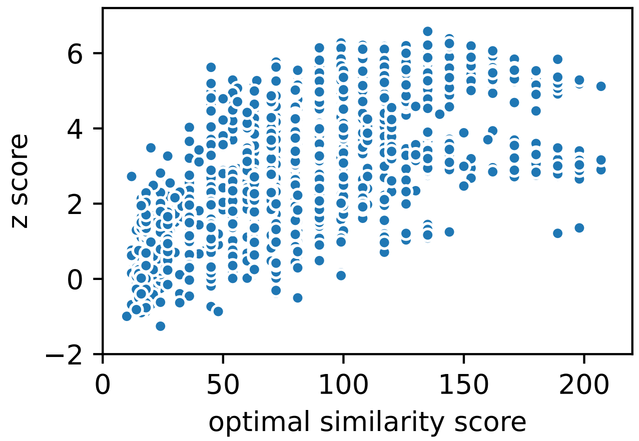


Figure S5: **Z score vs. optimal sequence similarity score for 10x genomics dataset. [3,4]**

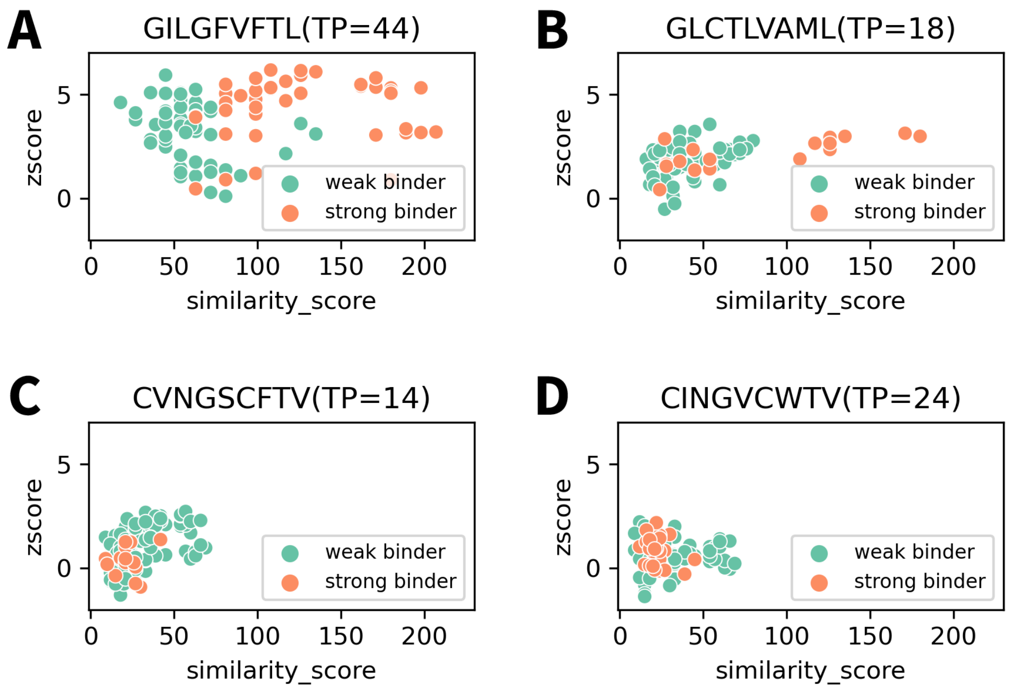


Figure S6: **Z score vs. optimal sequence similarity score for dataset from Grant *et al.* [5].**

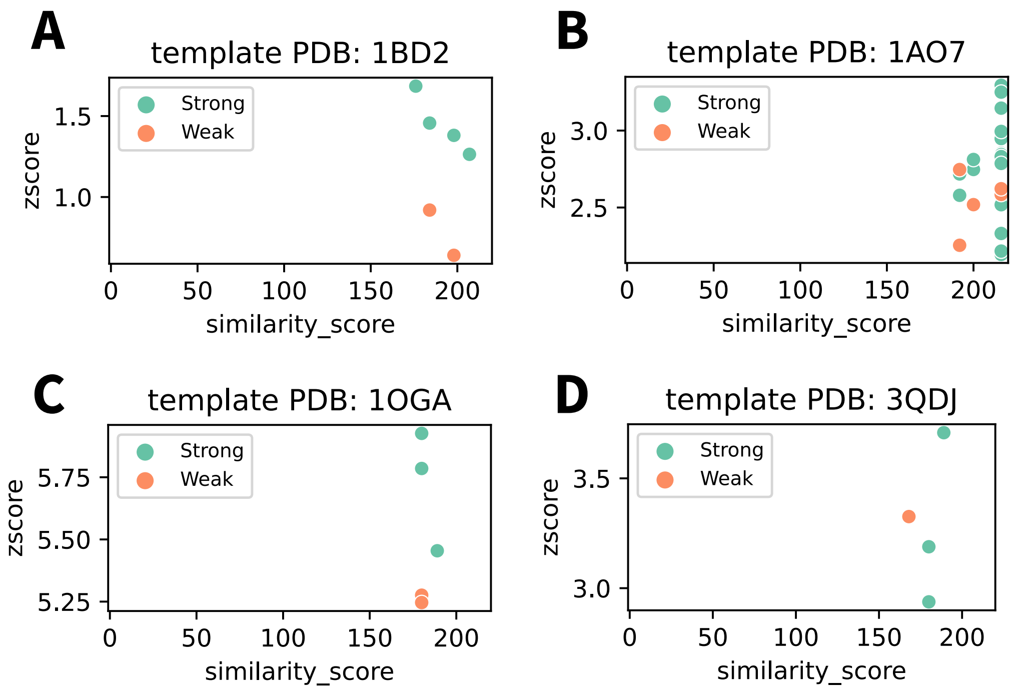


Figure S7: **Z score vs. optimal sequence similarity score for point-mutant weak binders in comparison with strong binders from ATLAS dataset [6].**
